# *ACPT* gene is inactivated in mammalian lineages that lack enamel and teeth

**DOI:** 10.1101/861633

**Authors:** Yuan Mu, Xin Huang, Rui Liu, Yulin Gai, Na Liang, Daiqing Yin, Lei Shan, Shixia Xu, Guang Yang

## Abstract

Loss of tooth or enamel is widespread in multiple mammal lineages. Although several studies have been reported, the evolutionary mechanisms of tooth / enamel loss are still unclear. Most previous studies have found that some tooth-related genes have been inactivated in toothless and / or enamel-less mammals, such as *ENAM*, *ODAM*, *C4orf26*, *AMBN*, *AMTN*, *DSPP*, etc. Here, we conducted evolutionary analyses on *ACPT* plays a key role in amelogenesis, to interrogate the mechanisms. We obtained the *ACPT* sequences from 116 species, including edentulous and enamel-less mammals, then evolutionary analyses were implemented. The results showed that variant ORF-disrupting mutations have been detected in *ACPT* coding region among nine edentulous baleen whales and three enamel-less taxa (pygmy sperm whale, aardvark, nine-banded armadillo). Furtherly, selective pressure uncovered that the selective constraints have been relaxed among all toothless and enamel-less lineages. Moreover, our results support the hypothesis that mineralized teeth were lost or degenerated in the common ancestor of crown Mysticeti through two shared single-base sites deletion in exon 4 and 5 of *ACPT* among all living baleen whales. *D*_N_ / *d*_S_ values on transitional branches were used to estimate *ACPT* inactivation times. In the case of aardvark, inactivation of *ACPT* was estimated at ~23.60-28.32 Ma, which is earlier than the oldest pangolin fossil time (*Orycteropus minutus*, ~19Ma), suggesting that *ACPT* inactivation may result in degeneration or loss of enamel. Conversely, the inactivation time of *ACPT* estimated in armadillo (~10.18-11.30 Ma) is later than the oldest fossil time, suggesting that inactivation of *ACPT* may result from degeneration or loss of enamel in these mammals. Our findings suggested that different mechanisms of degeneration of tooth / enamel might exist among toothless and enamel-less lineages during evolution. Our study further considered that *ACPT* is a novel gene for studying tooth evolution.

## Introduction

Dental innovations (such as differentiated dentitions and the evolution of tribosphenic molar, etc.) have been regarded as the great success of mammalian evolution and adaptation (Ungar 2010). Various sets of dentition facilitated mastication to gain energy from food efficiently, promoted major changes in feeding strategy, life history, and ecological niches for mammalian radiation during the past 220 million years ago (Ma) (Bergqvist 2003; Ungar 2010). However, in spite of their importance for animal survival, teeth have been lost independently in multiple mammal lineages, such as baleen whales and pangolins. In addition, some lineages have lost their outer enamel of teeth, such as pygmy sperm whale and dwarf sperm whale, aardvarks and species from Xenarthra (Davitbeal et al. 2009). Tooth loss and / or enamel loss is one of the most important field for mammalian tooth evolution.

Amelogenesis imperfecta (AI) and tooth loss are the diseases that characterized by genetic defects in the formation of enamel and teeth. Multiple studies have suggested these genetic disorders are mainly caused by mutations of protein-coding genes functioned in formation of enamel and teeth (Stephanopoulos et al. 2005; Smith et al. 2017). Of these genes, three enamel matrix protein genes (EMPs, i.e., *AMELX*, *AMBN* and *ENAM*), two enamel proteases genes (*MMP20* and *KLK4*), and some other related genes (e.g., *C4orf26*, *AMTN*, *ODAM*, *ACPT*, *DSPP*) have been confirmed to be candidate genes responsible for the diseases (Crawford et al. 2007; Smith et al. 2017). The variant inactivating mutations have been detected in these genes among toothless and enamel-less mammalian lineages. However, the mechanisms of tooth loss or enamel loss are still unclear.

It has been reported that *ACPT* was lower expressed in testicular cancer tissues compared to normal tissues and is regulated by steroid hormones (Yousef et al. 2001). Besides, *ACPT* is also expressed in the brain and acts as a tyrosine phosphatase to modulate signals mediated by ErbB4 (Fleisig et al. 2004). But, it is interesting to note that *ACPT* is expressed in secretory-stage ameloblasts (Seymen et al. 2016), which can induce odontoblasts differentiation, mineralization of dentin, and amelogenesis (Choi et al. 2016). Furthermore, there are some increasing evidences that homozygous missense variants of *ACPT* would lead to AI (e.g., c.226C>T, p.Arg76Cys; c.746C4T, p.P249L) (Seymen et al. 2016; Smith et al. 2017). Sharma et al. (2018) found that *ACPT* was pseudogene in minke whale (*Balaenoptera acutorostrata*), armadillo (*Dasypus novemcinctus*) and aardvark (*Orycteropus afer*), whose teeth or enamel are absent. These evidences suggested that *ACPT* should play an important role in amelogenesis.

To our knowledge, all extant Mysticeti, descended from toothed ancestors, have no teeth where instead of baleen (Uhen 2010). Paleontological evidences have shown that mineralized teeth were lost in the common ancestor of crown Mysticeti. Moreover, a transitional stage from tooth to baleen in stem mysticetes have been revealed in some taxa bearing both teeth and baleen (Deméré et al. 2008). Although many tooth-related genes have been revealed to be inactivated in various living mysticetes (e.g., *AMBN*, *ENAM*, *AMEL*, *AMTN*, *MMP20*, *C4orf26* and *DSPP*) (Deméré et al. 2008; Meredith et al. 2009; Meredith et al. 2011; Gasse et al. 2012; Delsuc et al. 2015; Springer et al. 2016; Springer et al. 2019), only the *MMP20* are commonly inactivated across all the living baleen whales, which indicates enamel common ancestor of baleen whales had already lost teeth (Meredith et al. 2011). This molecular evidence is consistent with earlier studies of paleontology and anatomy.

Despite its significance in mammalian enamel maturation, very little is known about *ACPT* evolutionary trajectory, relationship and function in mammals. To address this issue, we carried out a series of evolutionary analyses on *ACPT*, aim to uncover the evolutionary pattern of *ACPT* gene among mammalian lineages.

## Methods

### Sequences mining and BLAST searches

The full-length coding sequences (CDS) of *ACPT* gene were extracted from the OrthoMaM v10b (http://orthomam2.mbb.univ-montp2.fr/OrthoMaM_v10b10/), ENSEMBL (http://www.ensembl.org/index.html?redirect=no) and NCBI (http://www.ensembl.org/index.html?redirect=no) databases (Appendix Table 1). *ACPT* of some whales were extracted from their Genome and SRA database of NCBI (Appendix Table 2, 3). To further ensure the sites of inactivating mutation of toothless / enamel-less lineages, we used the CDSs of some representative placental species with well-annotated genomes (*Homo sapiens* [human], *Canis lupus familiaris* [Dog], *Bos taurus* [Cow], *Echinops telfairi* [Lesser hedgehog tenrec]) as queries including ∼50bp of flanking sequence on each exon. These sequences were used as queries to BLAST against toothless / enamel-less mammals from the closely related taxa to confirm the related inactivating mutation among baleen whales.

### Identification of inactivating mutations and functional sites and domains

The intact *ACPT* sequences (human, cow, tenrec) were used for identifying inactivating mutations (including mutation of initiation codons, frame-shift insertions and deletions, premature stop codons, splice sites mutation of intron / exon boundary [GT/AG], etc.). The inactivating mutation was identified based on BLAST searches against whole genomes of the relevant taxon from NCBI.

### Alignment and phylogenetic analysis of mammalian *ACPT*

The 116 mammalian *ACPT* sequences were aligned based on their amino acid translations using online PRANK (https://www.ebi.ac.uk/goldman-srv/webprank/), and then deleted the gaps and non-homologous regions by using GBLOCK, then we corrected the multiple sequences alignment (MSA) in MEGA 7 (Kumar et al. 2016) by eye.

A gene tree was reconstructed by Mrbayes 3.2 (Ronquist et al. 2012) with a general time reversible (GTR) substitution model and rate heterogeneity modeled with a Gamma distribution, as conducted by MrModeltest version 2 using the Akaike information criterion (AIC) (Nylander 2004). In bayesian analysis, four simultaneous runs with four chains each were run for two million generations, sampling every 1000 trees. The first 25% of these trees were discarded as burn-in when computing the consensus tree (50% majority rule).

### Selection analyses

To evaluate the selective pressure of relevant branches leading to enamel-less and toothless lineages respectively, we implemented *two ratio branch model* to calculate the ratio of the nonsynonymous substitution rate (*d*_N_) to the synonymous substitution rate (*d*_S_) (ω = *d*_N_/*d*_S_) by running CodeML in PAML 4.8a package (Yang 2007). We also recoded premature stop codons as missing data. Akaike information criterion (AIC) scores were used to select the most appropriate codon frequency model in CodeML. The *ACPT* gene tree exhibits different topological relationship compared to species tree, which may be unrelated to incomplete lineage sorting. In order to illuminate the detected signal reasonably and accurately, we used a species tree supported by some previous studies (Appendix Figure 1).

Refer to the methods of Springer and Gatesy (Springer and Gatesy 2018), several different branch categories have been considered during selective analyses: (1) One category accounted for ‘background’ branches, which are lineages with intact teeth and an intact copy of *ACPT*. (2) Nine branch categories to terminal branches with unique inactivating mutations (baleen whales), which lacks teeth. (3) Three branch categories to terminal branches with unique inactivating mutations (pygmy sperm whale, nine-banded armadillo and aardvark), whose enamel has been vestigial. (4) One branch categories were assigned for stem Mysticeti where mineralized teeth were degraded. (5) One branch categories were assigned for crown Mysticeti.

To better understand the selective pressure, a series of evolutionary models were compared in the likelihood. We first use the M0 model (Model A), which assumed that all branches in the phylogenetic tree has a common value, and compare it with the null hypothesis (Model B), which assumed that the common value in the phylogenetic tree is 1. To further understand whether the selective pressure on the lineages leading to pseudogenes was relaxed, we constructed Model C, which assumed that the branches with pseudogene had their own selection pressure ω_2_, while the background branches without pseudogenation was ω_1_, and then compared Model C with Model A. To further confirm whether the selective pressure on the lineages leading to pseudogenes was completely relaxed, we build the Model D, which assumed that the branches with pseudogene had their own selection pressure ω_2_ = 1, while the selective pressure of background branches was ω_1_, and then compared Model C with Model D.

### Estimation of inactivation times

To estimate when *ACPT* was inactivated in different lineages of Placentalia, we followed the procedure described in (Sharma et al. 2018). For a branch along which the gene was inactivated, this method assumes that a gene evolves under a selective pressure similar to that in other species until it is inactivated. Afterward, the gene is assumed to accumulate both synonymous and nonsynonymous mutations at a neutral rate. The K_a_ / K_s_ (K) value estimated for this entire branch is then the average of the K_a_ / K_s_ value for the part of the branch where the gene was under selection (K_s_), and the K_a_ / K_s_ value for the part of the branch where the gene evolved neutrally (K_n_ = 1). It is weighted by the proportion of time for which the gene was evolving under selection (T_s_ / T) and neutrally (T_n_ / T):

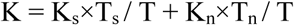

where T represents the time since the split from the last common ancestor. Using the lower and upper bound of the confidence interval for the species divergence time T obtained from TimeTree (http://www.timetree.org/) and using the K_a_/K_s_ value for mammals with a functional *ACPT*, one can estimate a lower and upper bound for T_n_ as:

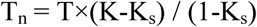

which provides an estimate of how long *ACPT* has been evolving neutrally.

## Results

### Characteriztion of *ACPT* sequence

The complete protein-coding sequence of *ACPT* in 116 taxon were used to alignment by PRANK. One or more inactivating mutations (frame-shift mutation, initial codon mutation, premature stop codons, splice site mutations, etc.) were detected in all placental taxa without teeth or without enamel (Figure 1, Appendix Table 4, Appendix Figure 2). For example, among toothless baleen whales, the initial codon mutation (n. ATG→GTG, p. M→V) was found in *Balaenoptera borealis*, *B. physalus*, *B. musculus*, *Eschrichtius robustus*, *Eubalaena glacialis*. Meanwhile, premature stop codons were found in *B. acutorostrata* and *B. bonaerensis*, frameshift indels were also found in baleen whales. Interestingly, two shared single-base site deletion was found on exon 4 and 5 of *ACPT* among all living baleen whales (Figure 1, Appendix Figure 2). The splice site mutations were detected in *B. edeni*, *B. omurai* and *Megaptera novaeangliae* (Appendix Figure 2). On the contrary, the premature stop codons were found in enamel-less *Dasypus novemcinctus* and *Orycteropus afer*. Besides, frameshift indels were found in *Kogia breviceps*.

**Figure 1.**
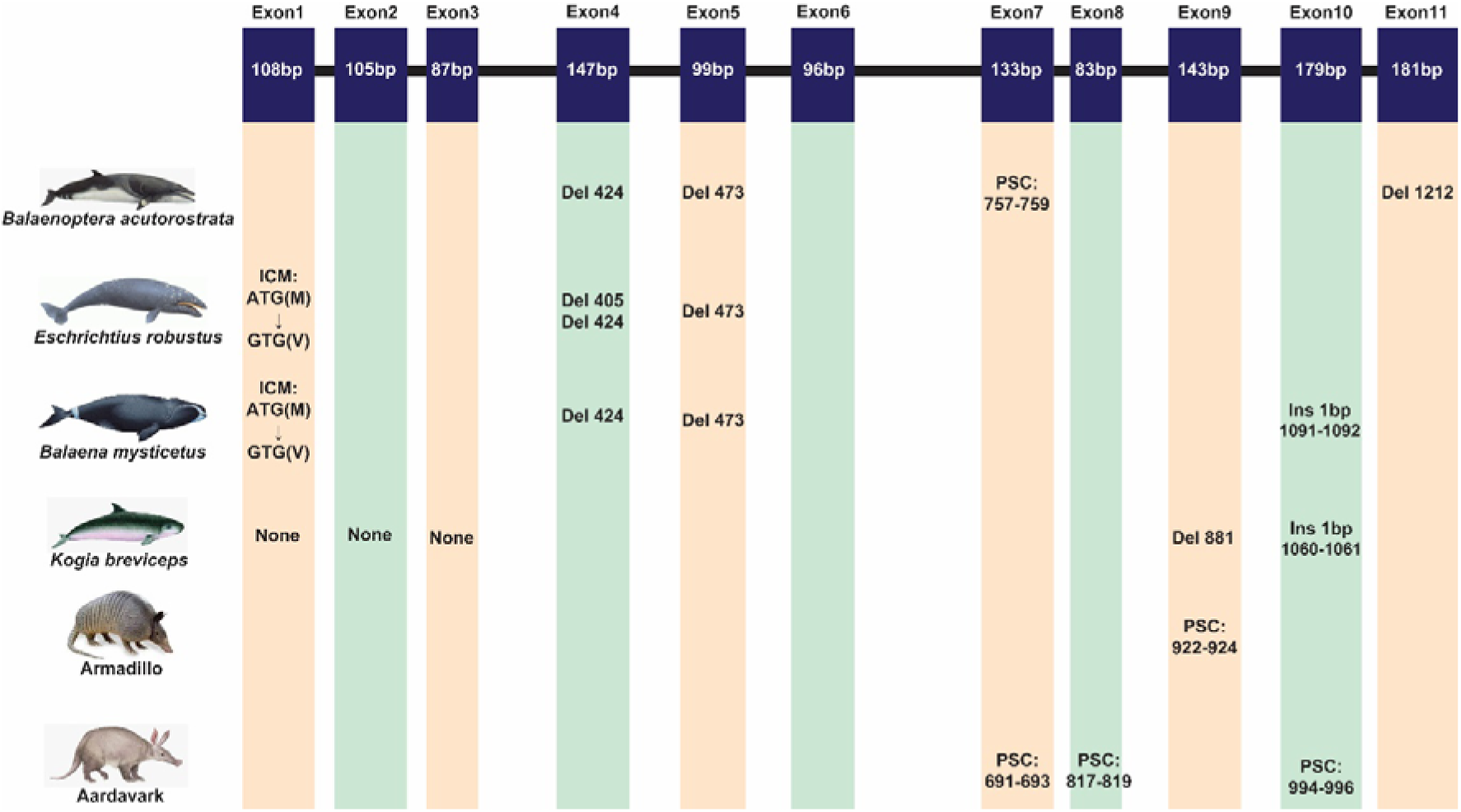
The inactivating mutation of *ACPT* gene in toothless/enamel-less mammals. (Abbreviation: ICM, initiation codon mutation; Del, deletion; Ins, insertion; PSC, premature stop codon)

Except for the species mentioned above, *ACPT* gene in other species were found to be intact. Nevertheless, some crucial amino acids mutation was found in toothed species, such as site 76 has been mutated (R76C) in *Neophocaena asiaeorientalis*..

### Reconstruction of *ACPT* gene tree

We recovered the *ACPT* gene tree with well-supported values by using Mrbayes method (Figure 2). In this gene tree, most of orders have been well reconstructed, and have high support rate, e.g., Cetartiodactyla, Perissodactyla, Eulipotyphla, Carnivora, Chiroptera etc. In addition, phylogenetic relationships of higher levels have also been well reconstructed, such as Laurasiatheria, Euarchontoglires, Boreoeutheria and Afrotheria. In this gene tree, bayesian posterior probability (PP) values of nearly 70% nodes are generally greater than 0.70. However, the relationship between some order level were relatively chaotic, such as Lagomorpha didn’t cluster with Rodentia, but as the sister group of Primate; Chiroptera and Carnivora clustered together first, and then they became sister group of Perissodactyla.

**Figure 2.**
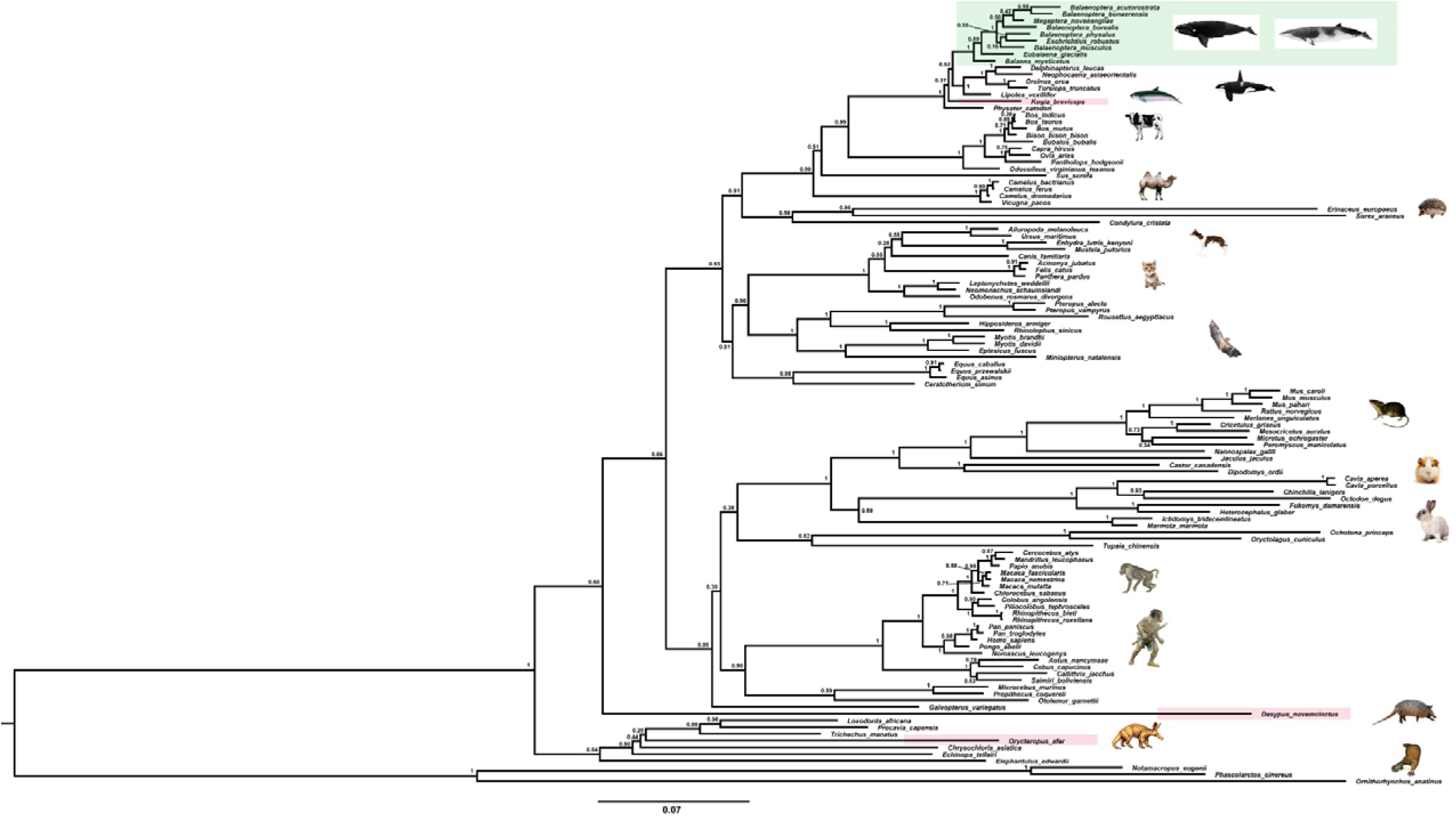
The BI phylogenetic relationship of 115 mammalian *ACPT* gene. (Nucleotide optimal substitution model: GTR+GAMMA; green box indicates toothless taxa, red boxes indicate enamel-less taxa)

### Evolutionary analyses among toothless and enamel-less mammals

We carried out the PAML analysis to detect the selective pressure of toothless / enamel-less lineages, and found the selective pressure of these toothless / enamel-less lineages (including ancestral nodes, terminal branches and even the whole toothless / enamel-less group) was significantly higher than that of background branches. For example, the terminal branch of *Balaenoptera physalus*: ω_1_=0.116, ω_2_=1.883; the terminal branch of *Megaptera novaeangliae*: ω_1_=0.116, ω_2_=0.641; the terminal branch of *Eschrichtius robustus*: ω_1_=0.116,ω_2_=2.688; the terminal branch of *Eubalaena glacialis*: ω_1_=0.116, ω_2_=0.503. A similar tendency was found in the terminal branches of other baleen whales, and further model comparison showed that the selective pressure of these branches had been completely relaxed. Whilst, much higher selective pressure was detected in the ancestral branch of stem mysticeti (ω_1_=0.120, ω_2_=0.436), even the clade of crown mysticeti (ω_1_=0.116, ω_2_=0.522). Meanwhile, higher selective pressure was detected among enamel-less lineages, such as the terminal branch of *Dasypus novemcinctus* (ω_1_=0.116, ω_2_=0.206), the terminal branch of *Orycteropus afer* (ω_1_=0.116, ω_2_=0.414), and the terminal branch of *Kogia breviceps* (ω_1_=0.116, ω_2_=0.581). And the selective pressure of these branches had been completely relaxed, except for the terminal branch of *K. breviceps* (Table S5).

### *ACPT* inactivation dates

Estimates of inactivation times for *ACPT* based on *d*_N_ / *d*_S_ ratios and equations in Sharma et al. (Sharma et al. 2018). The mean estimate for the inactivating time of *ACPT* on the branch of *K. breviceps*, *D. novemcinctus* and *O. afer* is 12.20-15.52Ma, 10.18-11.30Ma and 23.60-28.32Ma, respectively (Figure 3). The mean estimate for the inactivation of *ACPT* on the Mysticeti clade is 14.05-16.30Ma.

**Figure 3.**
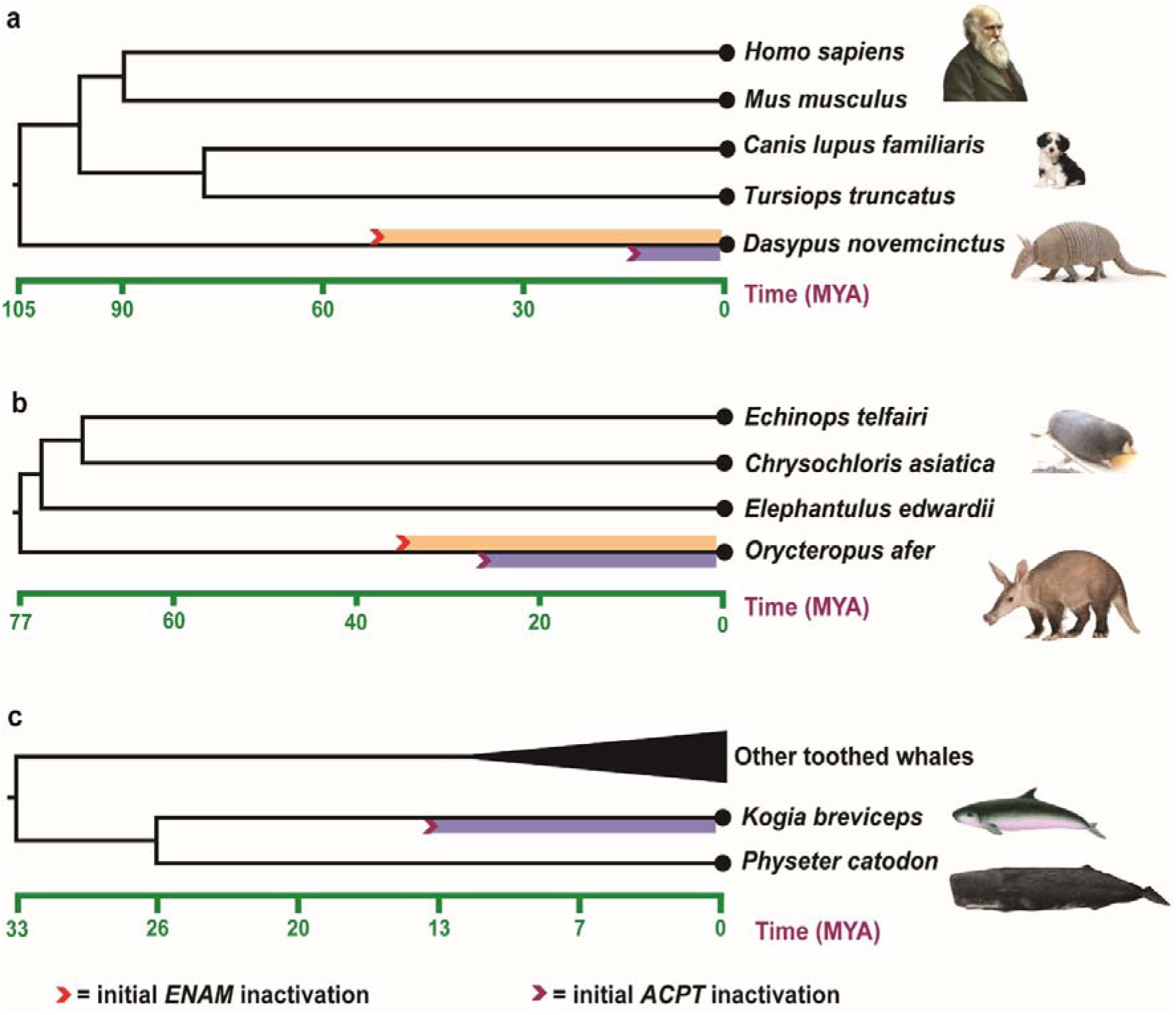
Estimated inactivation times of *ACPT* versus *ENAM*. (a) *Dasypus novemcinctus* (nine-banded armadillo), (b) *Orycteropus afer* (aardvark), (c) *Kogia breviceps* (pygmy sperm whale). The inactivation times of *ENAM* is from (Meredith, et al. 2009; Springer, et al. 2019).

## Discussion

### *ACPT* is a novel candidate gene for studying mammalian tooth loss and enamel loss

The well-conserved gene structure indicates that this organization was present in the last common mammalian ancestor, and the protein function was already defined 220 Ma (Madsen 2009). In our study, the number of *ACPT* exons are 11 in placental mammals, which encode 427 amino acids (human ACPT sequence as the reference sequence). Our study highlighted that four residues (191N, 269N, 330N and 339N) of the extracellular region were for glycosylation, two residues (41H and 289D) directly involved in catalysis. In addition, mutation in seven residues were reported that were responsible for AI (Seymen et al. 2016; Smith et al. 2017) (Appendix Figure 3). Three disulfide bond regions were identified, namely, site 159 to 378, site 214 to 312, site 353 to 357. In fact, we detected not only teratogenic mutations but also inactivated mutations in these functional sites and domains. For example, enamel in finless porpoise were degenerated (Ishiyama 1987), mutation in site 76 (R→C) was found in *N. asiaeorientalis*. Previous research has confirmed that site 76 mutated into Cys (C) in human ACPT would lead to hypoplastic AI (Seymen et al. 2016), from which this result further supported that teeth in finless porpoise were degenerated in molecular level. Of cause, some mutations were also detected in toothless and enamel-less lineages. The most obvious characteristics of *ACPT* is that different types of inactivating mutations were found in toothless and enamel-less mammals. Therefore, *ACPT* could be a candidate gene for AI and should be regarded as a target gene in the diagnosis of this genetic disease.

### Degeneration or loss of mineralized teeth in LCA of Mysticeti

Fossil evidence showed that the earliest ancestors of baleen whales possessed complete dentitions without baleen (such as *Janjucetus* and *Mammalodon*), and then evolved the baleen with teeth (such as *Aetiocetus*), until the lineages only baleen existed (e.g., *Eomysticetus* and *Micromysticetus*) (Fitzgerald 2006; Fitzgerald 2010). This supported the hypothesis that mineralized teeth were lost or degenerated in the common ancestor of crown Mysticeti. The fact is all living baleen whales lack teeth and instead baleen (Uhen 2010). However, the successive steps of vestigial tooth development was found in the fetal period of living baleen whales (Davitbeal et al. 2009; Thewissen 2018). Molecular sequences of some specific genes, such as *AMBN*, *ENAM*, *AMELX*, *AMTN*, *C4orf26* and *ODAM*, contain different types of inactivating mutations (e.g., stop codons, frameshift mutations, splice site mutations, etc.) in various mysticete species (Deméré et al. 2008; Meredith et al. 2009; Alhashimi et al. 2010; Gasse et al. 2012; Meredith et al. 2013; Delsuc et al. 2015; Springer et al. 2019), which is consistent with loss-of-teeth. But none of the inactivating mutations are shared by all living mysticetes species. Meredith et al. found a common insertion of CHR-2 SINE retroposon in *MMP20* gene among all living baleen whales (Meredith et al. 2011), which further confirmed the hypothesis that mineralized teeth were lost or degenerated in the common ancestor of crown Mysticeti in the molecular level. It has been confirmed that mutations or deletions of *MMP20* gene would result in thin and brittle enamel layer (Caterina et al. 2002). Some studies have confirmed that *ACPT* gene is responsible for the development of enamel, and mutations can also lead to amelogenesis imperfecta (Choi et al. 2016; Seymen et al. 2016; Smith et al. 2017). Different inactivating mutations was detected among all mysticete species in *ACPT* gene in our study, besides, two shared single-base sites deletion were found on exon 4 and 5 of *ACPT* among all living baleen whales, which result in loss of function. Similar to the result of Meredith et al. (Meredith et al. 2011), our study supported the hypothesis that that mineralized teeth were lost or degenerated in the common ancestor of crown Mysticeti.

### Is inactivation of *ACPT* neutral or adaptive?

The degeneration and / or loss of some morphological structures (such as limbs, teeth, and eyes, etc.) is a complex process that may result from the relaxation of the negative selection (neutral evolution), adaptive evolution (direct natural / positive selection to conserve energy and / or eliminate the disadvantageous effects of morphological structure), and / or gene pleiotropy (indirect selection on another traits) (Wang et al. 2006; Zhang 2008; Krishnan and Rohner 2017). Sharma et al. revealed that evolutionary gene losses are not only a consequence, but may also be causally involved in phenotypic adaptations (Sharma et al. 2018). By estimating the inactivation time of pseudogenes, and comparing with fossil evidence, we will speculate whether gene inactivation is due to the adaptive or neutral selection.

The record of enamel-degenerated armadillo fossil is significantly earlier than the estimated time of *ACPT* inactivation (10.18-11.30Ma) (Ciancio et al. 2014), which suggested gene loss as a consequence of adaptation is likely the result of the relaxation of the negative selection. The results further supported the previous study (Sharma et al. 2018). Whereas, the inactivation time of *ENAM* (~45.5Ma) and *ODAM* (~40.43 Ma, range 36.38-45.45Ma) is much older than inactivation date for *ACPT* in armadillo (Springer et al. 2019). Conversely, the inactivation time of *ENAM* is relatively earlier than the fossil record time (~3.5-6.5Ma). The inactivation of *ENAM* gene might be the one of most main causes of degeneration / loss of tooth enamel in armadillos.

Inactivation date for *ACPT* (23.60-28.32Ma) is relatively younger than inactivation dates for *ENAM* (28.8-35.3Ma) and *ODAM* (~30.7Ma) in *O. afer* (Meredith et al. 2009; Springer et al. 2019). However, estimates for *ACPT*, *ODAM* and *ENAM* inactivation are both older than the oldest fossil aardvark, *O. minutus*, which is ~19Ma (Patterson 1975). It strongly suggested that gene loss may be the reason, not the consequence, for degeneration and / or loss of enamel, which is different from the result of Sharma et al. (Sharma et al. 2018).

Cetacean includes both toothless Mysticeti and enamel-less *Kogia*. Relaxation of selective pressure was detected in both crown and stem Mysticeti (Appendix Table 5), which is consistent with the archaic toothless mysticete, e.g., *Eomysticetus whitmorei* (Deméré et al. 2008). Molecular evidence showed *ACPT* has been lost its function in LCA of Mysticeti. However, The inactivation time of *ACPT* in Mysticeti is 14.05-16.30Ma, which is much younger than the toothless mysticete (~30Ma) and the split of Mysticeti (~25.9Ma). Obviously, this is not consistent with the facts. It might be associated with relatively lower rates of frameshift accumulation during evolution of mysticete pseudogenes and long lifespan of mysticete (Meredith et al. 2009; Meredith et al. 2011). To our knowledge, whether adaptive or neutral, the shared single-base site deletion in *ACPT* fills an important gap in our understanding of the macroevolutionary transition leading from the LCA of crown Cetacean to the LCA of crown Mysticeti. Stem physeteroids (sperm whales) are known from the Miocene and had teeth with enamel (Bianucci and Landini 2010). Our results provide support for loss of the intact ACPT in *K. breviceps*. ACPT was reported that play key roles in amelogenesis and differentiation of odontoblasts (Choi et al. 2016; Seymen et al. 2016; Smith et al. 2017). Our result is in line with the enamel-less morphological structure in *K. breviceps*.

## Appendix files

**Appendix Figure 1** The tree topology used to conduct the selective pressure analysis in PAML.

**Appendix Figure 2** The detailed information about inactivated mutation of *ACPT* among relative cetaceans. (Orange represents initiation codon mutation, yellow represents deletion, green represents insertion, and blue represents two common deletion sites among all baleen whales).

**Appendix Figure 3** The information of mutation sites about amelogenesis imperfecta in ACPT protein sequence.

**Appendix Table 1** Mammalian species used in this study and the sources of sequences.

**Appendix Tables 2** The genome information of cetacean species used in this study.

**Appendix Table 3** SRA information of 4 baleen whales species used in this study.

**Appendix Table 4** The information of exon/intron boundary in relative whales (obtained by BLAST by using python script *in silico*)

**Appendix Table 5** Likelihood and omega values estimated under two ratio branch model on *ACPT* gene among toothless and enamel-less branches.

## Abbreviations

ACPT: Acid Phosphatase, Testicular
EMP: enamel matrix protein
PAML: phylogenetic analysis by maximum likelihood
PP: posterior probability
Ma: million years ago

## Declarations

### Ethics approval and consent to participate

Not applicable.

### Consent to publish

Not applicable.

### Availability of data and materials

The data generated and analyzed during this study are included in this article and its appendix files, including 5 tables and 6 figures.

### Competing interests

The authors declare that they have no competing interests.

### Funding

This work was financially supported by the National Key Program of Research and Development, Ministry of Science and Technology of China (grant no. 2016YFC0503200 to S.X.), the National Natural Science Foundation of China (NSFC) (grant nos. 31570379, 31772448 to S.X.) and the Priority Academic Program Development of Jiangsu Higher Education Institutions (PAPD). These funding bodies played no role in study design, data collection, analysis, interpretation or manuscript preparation.

### Authors’ contributions

Yuan Mu: contributed to conception and design, acquisition, analysis, and interpretation, drafted manuscript, critically revised manuscript, agrees to be accountable for all aspects of work ensuring integrity and accuracy.

Xin Huang: contributed to acquisition and analysis, drafted manuscript

Rui Liu: contributed to analysis and interpretation

Yulin Gai: contributed to design, critically revised manuscript

Na Liang: contributed to analysis, drafted manuscript

Daiqing Yin: contributed to conception, contributed to interpretation

Lei Shan: contributed to conception and design, critically revised manuscript

Shixia Xu: contributed to conception and design, interpretation, critically revised manuscript, gave final approval

Guang Yang: contributed to conception and design, critically revised manuscript, gave final approval, agrees to be accountable for all aspects of work ensuring integrity and accuracy

## Acknowledgements

We thank members of the Jiangsu Key Laboratory for Biodiversity and Biotechnology, Nanjing Normal University, for their contributions to this paper. The authors thank Mr. Xinrong Xu, Dr. Di Sun and Dr. Ran Tian, Dr. Zepeng Zhang, Dr. Simin Chai and Dr. Zhenpeng Yu for some helpful discussion. Special thanks to Dr. Zhengfei Wang for technical supports.

## References

Alhashimi N, Lafont AG, Delgado S, Kawasaki K, Sire JY. 2010. The enamelin genes in lizard, crocodile, and frog and the pseudogene in the chicken provide new insights on enamelin evolution in tetrapods. Mol Biol Evol. 27:2078–2094.

Bergqvist LP. 2003. The role of teeth in mammal history. Braz J Oral Sci. 2:249–257.

Bianucci G, Landini W. 2010. Killer sperm whale: a new basal physeteroid (Mammalia, Cetacea) from the Late Miocene of Italy. Zool J Linn Soc-Lond. 148:103–131.

Caterina JJ, Skobe Z, Shi J, Ding Y, Simmer JP, Birkedalhansen H, Bartlett JD. 2002. Enamelysin (MMP-20) deficient mice display an amelogenesis imperfecta phenotype. J Biol Chem. 277:49598–49604.

Choi H, Kim TH, Yun CY, Kim JW, Cho ES. 2016. Testicular acid phosphatase induces odontoblast differentiation and mineralization. Cell Tissue Res. 364:95–103.

Ciancio MR, Vieytes EC, Carlini AA. 2014. When xenarthrans had enamel: insights on the evolution of their hypsodonty and paleontological support for independent evolution in armadillos. Naturwissenschaften. 101:715–725.

Crawford PJ, Aldred M, Bloch-Zupan A. 2007. Amelogenesis imperfecta. Orphanet J Rare Dis. 2:17.

Davitbeal T, Tucker AS, Sire J. 2009. Loss of teeth and enamel in tetrapods: fossil record, genetic data and morphological adaptations. J Anat. 214:477–501.

Delsuc F, Gasse B, Sire JY. 2015. Evolutionary analysis of selective constraints identifies ameloblastin (AMBN) as a potential candidate for amelogenesis imperfecta. BMC Evol Biol. 15:148.

Deméré TA, Mcgowen MR, Berta A, Gatesy J. 2008. Morphological and molecular evidence for a stepwise evolutionary transition from teeth to baleen in mysticete whales. Syst Biol. 57:15–37.

Fitzgerald EM. 2006. A bizarre new toothed mysticete (Cetacea) from Australia and the early evolution of baleen whales. Proc R Soc B. 273:2955–2963.

Fitzgerald EMG. 2010. The morphology and systematics of Mammalodon colliveri (Cetacea: Mysticeti), a toothed mysticete from the Oligocene of Australia. Zool J Linn Soc-Lond. 158:367–476.

Fleisig H, El-Husseini ED, Vincent SR. 2004. Regulation of ErbB4 phosphorylation and cleavage by a novel histidine acid phosphatase. Neuroscience. 127:91–100.

Gasse B, Silvent J, Sire JY. 2012. Evolutionary analysis suggests that AMTN is enamel-specific and a candidate for AI. J Dent Res. 91:1085.

Ishiyama M. 1987. Enamel structure in odontocete whales. Scan Microsc. 1:1071–1079.

Krishnan J, Rohner N. 2017. Cavefish and the basis for eye loss. Phil Trans R Soc B. 372:20150487.

Kumar S, Stecher G, Tamura K. 2016. MEGA7: Molecular Evolutionary Genetics Analysis version 7.0 for bigger datasets. Mol Biol Evol. 33:1870–1874.

Madsen O. 2009. Mammals. In: The time tree of life. New York, NY: Oxford University Press.

Meredith RW, Gatesy J, Cheng J, Springer MS. 2011. Pseudogenization of the tooth gene enamelysin (MMP20) in the common ancestor of extant baleen whales. Proc R Soc B. 278:993–1002.

Meredith RW, Gatesy J, Murphy WJ, Ryder OA, Springer MS. 2009. Molecular decay of the tooth gene enamelin (ENAM) mirrors the loss of enamel in the fossil record of placental mammals. PLoS Genet. 5:e1000634.

Meredith RW, Gatesy J, Springer MS. 2013. Molecular decay of enamel matrix protein genes in turtles and other edentulous amniotes. BMC Evol Biol. 13:20.

Nylander JAA. 2004. MrModeltest v2. Program distributed by the author. Evolutionary Biology Centre, Uppsala University.

Patterson B. 1975. The fossil aardvarks (Mammalia: Tubulidentata). Bull Mus Comp Zool. 147:185–237.

Ronquist F, Teslenko M, Der Mark PV, Ayres DL, Darling AE, Hohna S, Larget B, Liu L, Suchard MA, Huelsenbeck JP. 2012. MrBayes 3.2: Efficient Bayesian Phylogenetic Inference and Model Choice across a Large Model Space. Syst Biol. 61:539–542.

Seymen F, Kim YJ, Lee YJ, Kang J, Kim TH, Choi H, Koruyucu M, Kasimoglu Y, Tuna EB, Gencay K. 2016. Recessive mutations in ACPT, encoding testicular acid phosphatase, cause hypoplastic amelogenesis imperfecta. Am J Hum Genet. 99:1199–1205.

Sharma V, Hecker N, Roscito JG, Foerster L, Langer BE, Hiller M. 2018. A genomics approach reveals insights into the importance of gene losses for mammalian adaptations. Nat Commun. 9:1215.

Smith CE, Whitehouse LL, Poulter JA, Brookes SJ, Day PF, Soldani F, Kirkham J, Inglehearn CF, Mighell AJ. 2017. Defects in the acid phosphatase ACPT cause recessive hypoplastic amelogenesis imperfecta. Eur J Hum Genet. 25:1015–1019.

Smith CEL, Poulter JA, Antanaviciute A, Kirkham J, Brookes SJ, Inglehearn CF, Mighell AJ. 2017. Amelogenesis imperfecta; genes, proteins, and pathways. Front Physiol. 8:435.

Springer M, Emerling C, Gatesy J, Randall J, Collin M, Hecker N, Hiller M, Delsuc F. 2019. Odontogenic ameloblast-associated (ODAM) is inactivated in toothless/enamelless placental mammals and toothed whales. BMC Evol Biol. 19:31.

Springer MS, Gatesy J. 2018. Evolution of the MC5R gene in placental mammals with evidence for its inactivation in multiple lineages that lack sebaceous glands. Mol Phylogenet Evol. 120:364–374.

Springer MS, Starrett J, Morin PA, Lanzetti A, Hayashi C, Gatesy J. 2016. Inactivation of C4orf26 in toothless placental mammals. Mol Phylogenet Evol. 95:34–45.

Stephanopoulos G, Garefalaki ME, Lyroudia K. 2005. Genes and related proteins involved in amelogenesis imperfecta. J Dent Res. 84:1117.

Thewissen JGM. 2018. Highlights of cetacean embryology. Aquat Mamm. 44:591–602.

Uhen MD. 2010. The Origin(s) of Whales. Annu Rev Earth Planet Sci. 38:189–219.

Ungar PS. 2010. Mammal teeth: origin, evolution, and diversity: Johns Hopkins University Press, Baltimore, MD.

Wang X, Grus WE, Zhang J. 2006. Gene losses during human origins. PLoS Biol. 4:e52.

Yang Z. 2007. PAML 4: Phylogenetic Analysis by Maximum Likelihood. Mol Biol Evol. 24:1586–1591.

Yousef GM, Diamandis M, Jung K, Diamandis EP. 2001. Molecular cloning of a novel human acid phosphatase gene (*ACPT*) that is highly expressed in the testis. Genomics. 74:385–395.

Zhang JZ. 2008. Positive selection, not negative selection, in the pseudogenization of rcsA in Yersinia pestis. P Natl Acad Sci USA. 105:E69.

