## Supplemental Tables and Figures for "*ACPT* gene is inactivated in mammalian lineages that lack enamel and teeth"

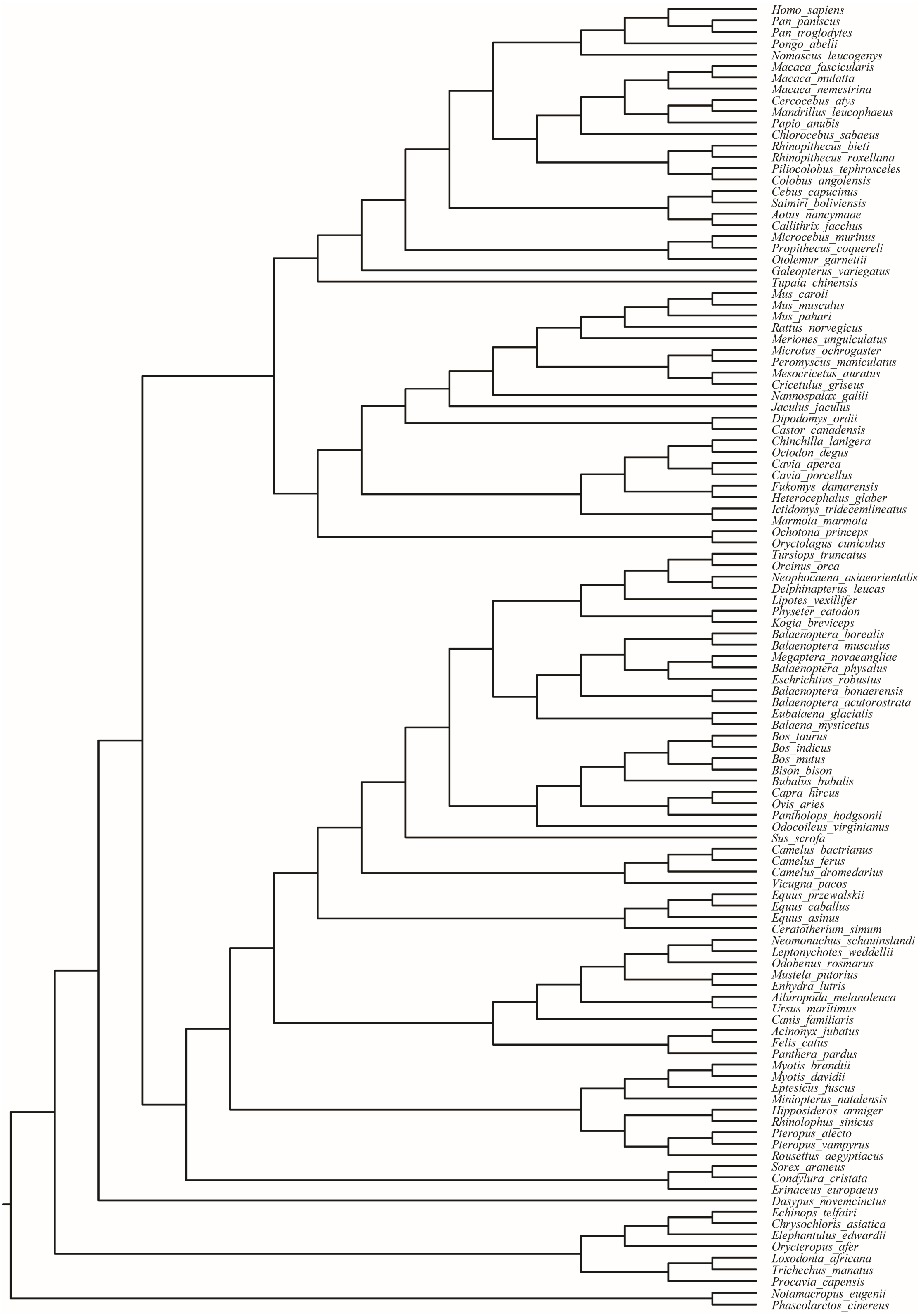


Appendix Figure 1 The tree topology of *ACPT* used to conduct the selective pressure analysis in PAML. The phylogeny is based on OrthoMaM (from: http://orthomam2.mbb.univ-montp2.fr/OrthoMaM_v10b10/) and some previous researches (Celine et al. 2019; Waddell et al. 2001; Sergey et al. 2007; Zhou et al. 2012; Gatesy et al. 2013; Kuntner et al. 2011)

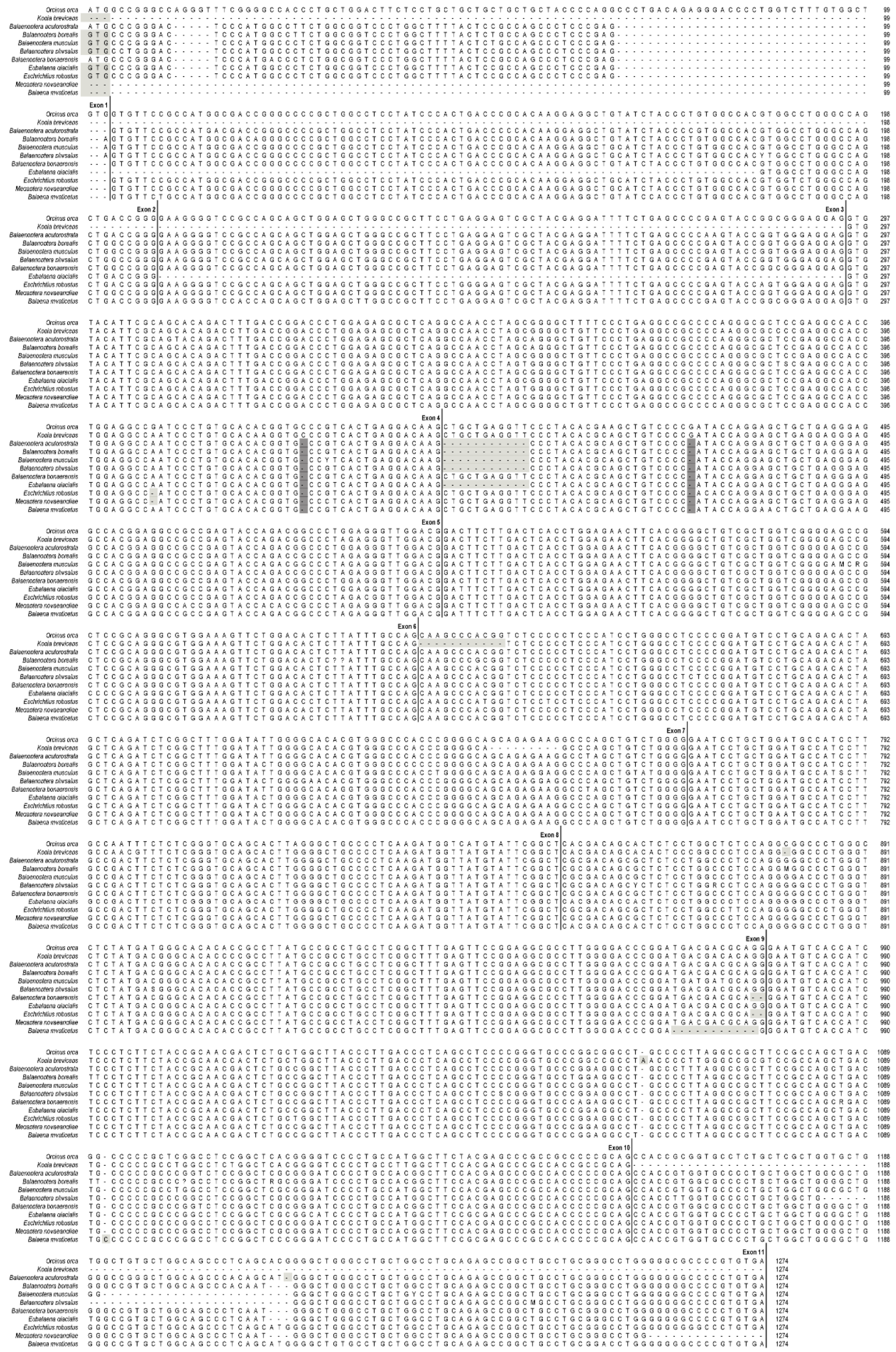

Appendix Figure 2 The detailed information about inactivated mutation of *ACPT* among relative cetaceans. (Light grey represents different type of inactivated mutation, initiation codon mutation, indels; dark grey represents two shared deletion sites in exon 4 and 5 among all baleen whales, respectively).

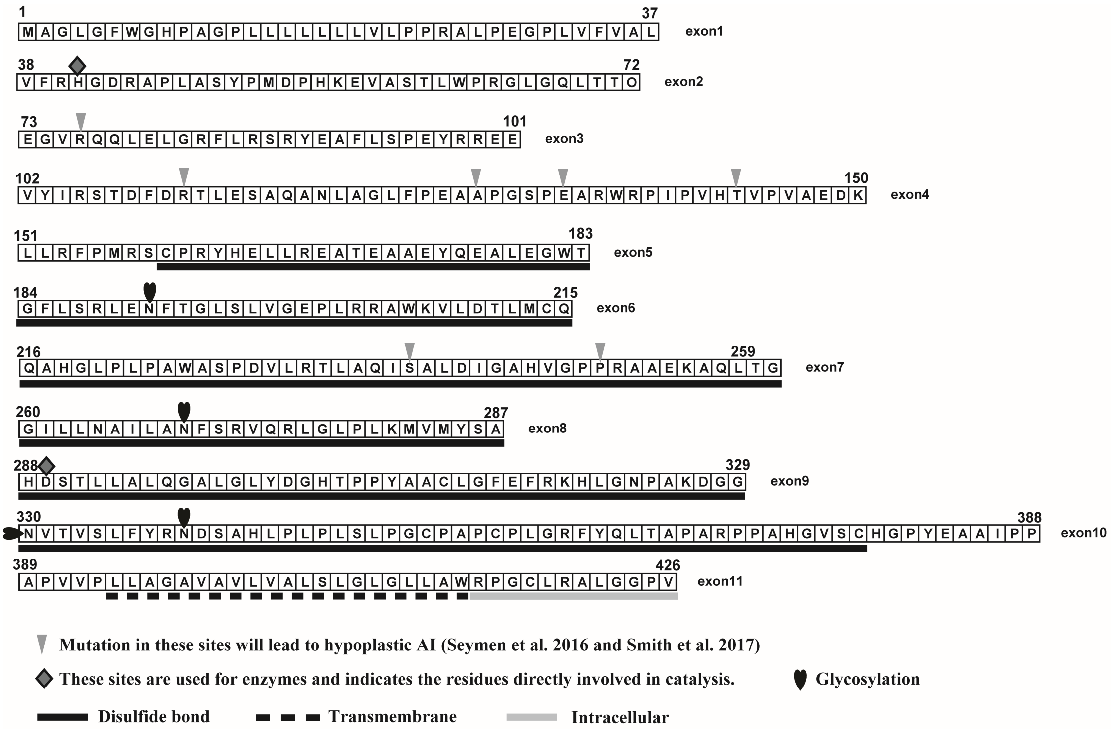


Appendix Figure 3 The information of mutation sites in ACPT protein sequence about amelogenesis imperfecta.

Appendix Table 1 Mammalian species used in this study and the sources of sequences.

| Species | *ACPT* | Species | *ACPT* | Species | *ACPT* |
|---|---|---|---|---|---|
| *Acinonyx_jubatus* | XM_027037126.1 | *Felis_catus* | ENSFCAG00000018727 | *Oryctolagus_cuniculus* | XM_017340604.1 |
| *Ailuropoda_melanoleuca* | ENSAMEG00000012861 | *Fukomys_damarensis* | ENSFDAG00000014874 | *Otolemur_garnettii* | XM_003801446.2 |
| *Aotus_nancymaae* | ENSANAG00000032261 | *Galeopterus_variegatus* | 103608037 | *Ovis_aries* | ENSOARG00000014152 |
| *Balaenoptera_acutorostrata* | XM_007179955.1 | *Gorilla_gorilla* | — | *Pan_paniscus* | ENSPPAG00000010927 |
| *Balaena_mysticetus* | bmy_14018 | *Heterocephalus_glaber* | ENSHGLG00000004949 | *Pan_troglodytes* | ENSPTRG00000011352 |
| *Bison_bison* | 104995692 | *Hipposideros_armiger* | 109390983 | *Panthera_pardus* | 109252997 |
| *Bos_taurus* | ENSBTAG00000015115 | *Homo_sapiens* | ENSG00000142513 | *Pantholops_hodgsonii* | XM_005955418.1 |
| *Bos_indicus* | 109571909 | *Ictidomys_tridecemlineatus* | ENSSTOG00000010097 | *Papio_anubis* | ENSPANG00000010665 |
| *Bos_mutus* | 102283597 | *Jaculus_jaculus* | ENSJJAG00000014058 | *Peromyscus_maniculatus* | ENSPEMG00000008909 |
| *Bubalus_bubalis* | 102407119 | *Lagenorhynchus_obliquidens* | — | *Phascolarctos_cinereus* | 110197749 |
| *Callithrix_jacchus* | ENSCJAG00000016684 | *Leptonychotes_weddellii* | 102742150 | *Phocoena_phocoena* | — |
| *Canis_familiaris* | ENSCAFG00000002920 | *Lipotes_vexillifer* | XM_007463413.1 | *Physeter_catodon* | 102977978 |

| | | | | | |
|---|---|---|---|---|---|
| *Cavia_aperea* | ENSCAPG00000001611 | *Loxodonta_africana* | XM_003406840.2 | *Piliocolobus_tephrosceles* | 111519955 |
| *Cavia_porcellus* | ENSCPOG00000026806 | *Macaca_fascicularis* | ENSMFAG00000003653 | *Pongo_abelii* | ENSPPYG00000010294 |
| *Camelus_bactrianus* | 105062672 | *Macaca_mulatta* | ENSMMUG00000003583 | *Procavia_capensis* | ENSPCAG00000003632 |
| *Camelus_dromedarius* | 105100227 | *Macaca_nemestrina* | ENSMNEG00000037245 | *Propithecus_coquereli* | ENSPCOG00000024911 |
| *Camelus_ferus* | 102516486 | *Mandrillus_leucophaeus* | ENSMLEG00000019431 | *Pteropus_alecto* | 102898726 |
| *Capra_hircus* | 102191826 | *Marmota_marmota* | 107151208 | *Pteropus_vampyrus* | ENSPVAG00000003432 |
| *Castor_canadensis* | 109683865 | *Meriones_unguiculatus* | 110541101 | *Rattus_norvegicus* | ENSRNOG00000021659 |
| *Ceratotherium_simum* | 101393590 | *Mesocricetus_auratus* | ENSMAUG00000016895 | *Rhinopithecus_bieti* | ENSRBIG00000027329 |
| *Cebus_capucinus* | ENSCCAG00000037802 | *Miniopterus_natalensis* | 107531832 | *Rhinopithecus_roxellana* | ENSRROG00000029516 |
| *Cercocebus_atys* | ENSCATG00000032166 | *Microcebus_murinus* | ENSMICG00000000521 | *Rhinolophus_sinicus* | 109452514 |
| *Chrysochloris_asiatica* | XM_006868138.1 | *Microtus_ochrogaster* | ENSMOCG00000022668 | *Rousettus_aegyptiacus* | 107500452 |
| *Chinchilla_lanigera* | ENSCLAG00000006023 | *Mus_caroli* | XM_021167373.1 | *Saimiri_boliviensis* | ENSSBOG00000026701 |
| *Chlorocebus_sabaeus* | ENSCSAG00000002075 | *Mus_musculus* | ENSMUSG00000012777 | *Sorex_araneus* | XM_004619766.1 |

| | | | | | |
|---|---|---|---|---|---|
| *Choloepus_hoffmanni* | — | *Mus_pahari* | XM_021222251.1 | *Sus_scrofa* | ENSSSCG00000024736 |
| *Colobus_angolensis* | ENSCANG00000025394 | *Mustela_putorius* | ENSMPUG00000001610 | *Trichechus_manatus* | 101344314 |
| *Condylura_cristata* | XM_004694037.1 | *Myotis_brandtii* | 102249381 | *Tupaia_chinensis* | 102500938 |
| *Cricetulus_griseus* | ENSCGRG00001003687 | *Myotis_davidii* | 102769794 | *Tursiops_truncatus* | ENSTTRG00000011047 |
| *Dasypus_novemcinctus* | XM_023585546.1 | *Myotis_lucifugus* | — | *Ursus_maritimus* | 103657347 |
| *Delphinapterus_leucas* | 111180510 | *Nannospalax_galili* | ENSNGAG00000020915 | *Vicugna_pacos* | ENSVPAG00000009708 |
| *Dipodomys_ordii* | ENSDORG00000013632 | *Neomonachus_schauinslandi* | 110572992 | *Orycteropus_afer* | XM_007957636.1 |
| *Echinops_telfairi* | XM_004710452.1 | *Neophocaena_asiaeorientalis* | XM_024768516.1 | *Equus_caballus* | XM_001917445.3 |
| *Elephantulus_edwardii* | 102863577 | *Nomascus_leucogenys* | ENSNLEG00000005747 | *Erinaceus_europaeus* | XM_007531289.1 |
| *Enhydra_lutris* | 111160580 | *Notamacropus_eugenii* | ENSMEUG00000008524 | *Eschrichtius_robustus* | Gonem-blast |
| *Eptesicus_fuscus* | 103297603 | *Ochotona_princeps* | ENSOPRG00000007351 | *Odocoileus_virginianus* | 110135451 |
| *Equus_asinus* | 106845328 | *Octodon_degus* | ENSODEG00000015849 | *Orcinus_orca* | 101283727 |
| *Equus_przewalskii* | 103540878 | *Odobenus_rosmarus* | 101374821 | | |

Appendix Table 2 The genome information of cetacean species used in this study.

| Species | Version | Coverage | Genebank Assembly Accession | Assembly method | Sequencing technology |
|---|---|---|---|---|---|
| *Balaena mysticetus* | Database Statistics (v1.0) | ~150x | Source: The Bowhead Whale Genome Resource (http://www.bowhead-whale.org/) | ALLPATHS-LG | Illumina HiSeq |
| *Balaenoptera bonaerensis* | ASM97880v1 | 60x | GCA_000978805.1 | PLATANUS v. 1.2.1 | Illumina HiSeq2000 |
| *Balaenoptera acutorostrata* | BalAcu1.0 | 92x | GCA_000493695.1 | SOAPdenovo v. 16-Mar-2012 | Illumina HiSeq 2000 |
| *Eschrichtius robustus* | EscRob_v1_BIUU | 40.5x | GCA_004363415.1 | DISCOVAR de novo v. discovardenovo-52488 | Illumina HiSeq |
| *Kogia breviceps* | KogBre_v1_BIUU | 38.8x | GCA_004363705.1 | DISCOVAR de novo v. discovardenovo-52488 | Illumina HiSeq |
| *Megaptera novaeangliae* | megNov1 | 102.0x | GCA_004329385.1 | Meraculous + HiRise v. Feb-2016 | Illumina HiSeq |

Appendix Table 3 The SRA information of 4 baleen whales species used in this study.

| **Species** | **Accession** | **Library name** | **Sample** | **Size** | **Relative information** |
|---|---|---|---|---|---|
| *Balaenoptera musculus* | SRX2901261 | Bmu_8065(Bmus_1) | cell culture | 45.3Gb | 1 ILLUMINA (Illumina HiSeq 2000) run: 559.5M spots, 113G bases, 45.3Gb downloads |
| *Balaenoptera borealis* | SRX2901260 | Bbo_E91(Bbor_1) | cell culture | 20.9Gb | 1 ILLUMINA (Illumina HiSeq 2500) run: 185.8M spots, 33.5G bases, 20.9Gb downloads |
| | SRX2901259 | Bbo_D27(Bbor_2) | cell culture | 20.5Gb | 1 ILLUMINA (Illumina HiSeq 2500) run: 184.5M spots, 33.2G bases, 20.5Gb downloads |
| *Eubalaena glacialis* | SRX2901265 | Egl_F68(Egla_1) | cell culture | 20.7Gb | 1 ILLUMINA (Illumina HiSeq 2500) run: 186M spots, 33.5G bases, 20.7Gb downloads |
| *Balaenoptera physalus* | SRX2901262 | Bph_4966_PE300_1 | cell culture | 17.1Gb | 1 ILLUMINA (Illumina HiSeq 2000) run: 205M spots, 41.4G bases, 17.1Gb downloads |

Appendix Table 4 The information of exon/intron boundary in relative whales (obtained by BLAST by using python script *in silico*)

| Species | Exon1 | Intron1 | Exon2 | Intron2 |
|---|---|---|---|---|
| *Balaena mysticetus* | …… | …...CTCCCCAG | GTGTTC……ACCGGG | GTGAGAAG……GTCCCCAG |
| *Balaenoptera bonaerensis* | …… | …...CTCCCCAG | GTGTTC……GCCGGG | GTGAGAAG……GTCCCCAG |
| *Balaenoptera acutorostrata* | …… | …...CTCCCCAG | GTGTTC…...ACCGGG | GTGAGAAG…...GTACCCAG |
| *Eubalaena japonica* | …… | …...CTCCCC**AC** | GTGTTC……ACCGGG | GTGAGAAG…...GTCCCCAG |
| *Eschrichtius robustus* | …… | …...CTCCCCAG | GTGTTC…...ACCGGG | GTGAGAAG…...GTCCCCAG |
| *Megaptera novaeangliae* | …… | …...CTCCCCAG | GTGTTC……GCCGGG | GTGAGAAG…...GTCCCCAG |
| *Kogia breviceps* | | | | |

Continued table

| Exon3 | Intron3 | Exon4 | Intron4 | Exon5 |
|---|---|---|---|---|
| GAAGGG……GAGGAG | GTACTGCC……GCACCCAG | GTGTAC…...GACAAG | GTCAGGGG…...TCTTCCAG | CTGCTG…...TGGACG |
| GAAGGG……GAGGAG | GTACTGCC……GCACCCAG | GTGTAC……GACAAG | GTCAGGGG……TCCTCCAG | CTGCTG……TGGACG |
| GAAGGG……GAGGAG | GTACTGCC……GCACCCAG | GTGTAC…...GACAAG | GTCAGGGG……NNNNNNNN | NNNNNN……TGGACG |
| GAAGGG……GAGGAG | GTACTGCC……GCACCCAG | GTGTAC……GACAAG | GTCAGGGG…...TCCTCCAG | CTGCTG……TGGACG |
| GAAGGG…...GAGGAG | GTACTGCC……GCACCCAG | GTGTAC……GACAAG | GTCAGGGG……TCCTCCAG | CTGCTG……TGGACG |
| GAAGGG……GAGGAG | GTACTGCC…...GCACCCAG | GTGTAC……GACAAG | GTCAGGGG…...TCCTCCAG | CTGCTG……TGGACG |
| | …….GCACCCAG | GTGTAC……GACAAG | GTCAGGGG…...TCCTCCAG | CTGCTG……TGGACG |

Continued table

| Intron5 | Exon6 | Intron6 | Exon7 | Intron7 |
|---|---|---|---|---|
| GTGAGCAG…...GCATCCAG | GATTTC…...TGCCAG | GTGGGTCC…...CTCCCCAG | CAAGCC…...CTGGGG | GTGAGGTG…...CCTGTCAG |
| GTGAGCAC……GCGTCCAG | GACTTC……TGCCAG | GTGGGTCC……CTCCCCAG | CAAGCC……CTGGGG | GTGAGGTG…...CCTGTCAG |
| GTGAGCAG…...GCGTCCAG | GACTTC…...TGCCAG | GTGGGTCC……CTCCCCAG | CAAGCC……CTGGGG | GTGAGGTG…...CCTGTCAG |
| GTGAGCAA……GCATCCAG | GATTTC……TGCCAG | GTGGGTCC…...CTCCCCAG | CAAGCC…...CTGGGG | GTGAGGTG……CCTGTCAG |
| GTGATCAG……GCGTCCAG | GACTTC…...TGCCAG | GTGGGTCC…...CTCCCCAG | CAAGCC……CTGGGG | GTGAGGTG…...CCTGTCAG |
| GTGAGCAG……GCGTCCAG | GACTTC…...TGCCAG | GTGGGTCC…...CTCCCCAG | CAAGCC……CTGGGG | GTGAGGTG……CCTGTCAG |
| GTGAGCGA…...GCATCCAG | GACTTC…...TGCCAG | GTGGCTCC……CCCCTTC... | ...TCTCCC…...CTGGGG | GTGAGGTG…...CCTGTCAG |

Continued table

| Exon8 | Intron8 | Exon9 | Intron9 | Exon10 |
|---|---|---|---|---|
| GAATCC…...TCGGCT | GTGAGTCT……GCCTGCAG | CACGAC…...CGCAGG | GTGAGGAG…...TCCTCCAG | GGATGT……CCGCAG |
| GAATCC…...TCGGCT | GTGAGTCT……GCCTGCAG | CGCGAC……GACGCA | GTGTGAGG…...CCCTCCAG | GGATGT……CCGCAG |
| GAATCC……TCGGCT | GTGAGTCT……GCCTGCAG | CGCGAC……GACGCA | GGGTGGTA…...CCCTCCAG | GGATGT…...CCGCAG |
| GAATCT……TCGGCT | GTGAGTCT……GCCTGCAA | CACGAC…...CGCAGG | GTGAGGAG…...TCCTACAG | GGATGT……CCGCAG |
| GAATCC…...TCGGCT | GTGAGTCT……GCCTGCAG | CGCGAC……GACGCA | GGGTGAGG…...CCCTCCAG | GGATGT…...CCGCAG |
| GAATCC……TCGGCT | GTGAGTCT……GCCCGCAG | CGCGAC…...CGCAGG | GTGAGGAG……CCTCCAAG | GATGTC……CCGCAG |
| GAATCC……TCGGCT | GTGAGTCT……GCCCGCAG | CACGAC……CACAGG | GTGAGGAG…...CCCTCCAG | GAATGT…...CCGCAG |

Continued table

| Intron10 | Exon11 |
|---|---|
| GTGACGGC…...CCCCCCAG | CCACCG…...CCCGTGTGA |
| …… | CCACCG……CCCCTGTGA |
| GTGACGGC…...CCCCGCAG | CCACCG…...CCCCTGTGA |
| GTGACGGC……CCCCCCAG | CCACCG…...CCCGTGTGA |
| GTGACGGC…...CCCTGCAG | CCACCG…..GCCCTGTGA |
| GTGACGGC…... | CCACCG……GCCTGG |
| GTGACGGC…... | |

NOTE: The red GT/AG is the normal boundary of intro/exon. The blue color represents the splice mutation.

Appendix Table 5 Likelihood and $\omega$ values estimated under two ratio branch model on *ACPT* gene among toothless and enamel-less branches.

| **Models and some relative branches** | **$\omega$** | **-ln L** | **np** | **Models comparison** | **2Δ (ln L)** | ***P*-value** |
|---|---|---|---|---|---|---|
| **The terminal branch of *Balaenoptera physalus*** | | | | | | |
| A. All branches have one $\omega$ | 0.118 | 23204.622 | 208 | | | |
| B. All branches have one $\omega = 1$ | 1 | 25437.563 | 207 | A vs B | 4465.882 | 0 |
| C. The terminal branch of *Balaenoptera physalus* with pseudogenized *ACPT* has $\omega_2$, others have $\omega_1$ | $\omega_1=0.116$ $\omega_2=1.883$ | 23190.070 | 209 | A vs C | 29.104 | <0.01 |
| D. The terminal branch of *Balaenoptera physalus* with pseudogenized *ACPT* has $\omega_2 = 1$, others have $\omega_1$ | $\omega_1=0.116$ $\omega_2=1$ | 23190.624 | 208 | D vs C | 1.108 | 0.293 |
| **The terminal branch of *Megaptera novaeangliae*** | | | | | | |
| A. All branches have one $\omega$ | 0.117 | 23161.978 | 208 | | | |
| B. All branches have one $\omega = 1$ | 1 | 25405.725 | 207 | A vs B | 4487.494 | 0 |
| C. The terminal branch of *Megaptera novaeangliae* with pseudogenized *ACPT* has $\omega_2$, others have $\omega_1$ | $\omega_1=0.116$ $\omega_2=0.641$ | 23156.277 | 209 | A vs C | 11.402 | <0.01 |
| D. The terminal branch of *Megaptera novaeangliae* with pseudogenized *ACPT* has $\omega_2 = 1$, others have $\omega_1$ | $\omega_1=0.116$ $\omega_2=1$ | 23156.626 | 208 | D vs C | 0.698 | 0.403 |
| **The terminal branch of *Balaena mysticetus*** | | | | | | |
| A. All branches have one $\omega$ | 0.117 | 23149.887 | 208 | | | |
| B. All branches have one $\omega = 1$ | 1 | 25394.078 | 207 | A vs B | 4488.382 | 0 |
| C. The terminal branch of *Balaena mysticetus* with pseudogenized | $\omega_1=0.116$ $\omega_2=0.551$ | 23146.023 | 209 | A vs C | 7.728 | <0.01 |

| | | | | | | | |
|---|---|---|---|---|---|---|---|
| *ACPT* has $\omega_2$, others have $\omega_1$ | | | | | | | |
| D. The terminal branch of *Balaena mysticetus* with pseudogenized *ACPT* has $\omega_2 = 1$, others have $\omega_1$ | $\omega_1$=0.116 | $\omega_2$=1 | 23146.532 | 208 | D vs C | 0.509 | 0.476 |
| **The terminal branch of *Eschrichtius robustus*** | | | | | | | |
| A. All branches have one $\omega$ | 0.117 | | 23181.416 | 208 | | | |
| B. All branches have one $\omega = 1$ | 1 | | 25415.085 | 207 | A vs B | 4467.338 | 0 |
| C. The terminal branch of *Eschrichtius robustus* with pseudogenized *ACPT* has $\omega_2$, others have $\omega_1$ | $\omega_1$=0.116 | $\omega_2$=2.688 | 23167.178 | 209 | A vs C | 14.238 | <0.01 |
| D. The terminal branch of *Eschrichtius robustus* with pseudogenized *ACPT* has $\omega_2 = 1$, others have $\omega_1$ | $\omega_1$=0.116 | $\omega_2$=1 | 23168.159 | 208 | D vs C | 1.962 | 0.161 |
| **The terminal branch of *Balaenoptera musculus*** | | | | | | | |
| A. All branches have one $\omega$ | 0.117 | | 23199.599 | 208 | | | |
| B. All branches have one $\omega = 1$ | 1 | | 25435.255 | 207 | A vs B | 4471.312 | 0 |
| C. The terminal branch of *Balaenoptera musculus* with pseudogenized *ACPT* has $\omega_2$, others have $\omega_1$ | $\omega_1$=0.116 | $\omega_2$=1.395 | 23187.344 | 209 | A vs C | 24.510 | <0.01 |
| D. The terminal branch of *Balaenoptera musculus* with pseudogenized *ACPT* has $\omega_2 = 1$, others have $\omega_1$ | $\omega_1$=0.116 | $\omega_2$=1 | 23188.136 | 208 | D vs C | 1.584 | 0.208 |
| **The terminal branch of *Eubalaena glacialis*** | | | | | | | |
| A. All branches have one $\omega$ | 0.117 | | 23163.917 | 208 | | | |
| B. All branches have one $\omega = 1$ | 1 | | 25407.391 | 207 | A vs B | 4486.948 | 0 |
| C. The terminal branch of *Eubalaena glacialis* with pseudogenized | $\omega_1$=0.116 | $\omega_2$=0.503 | 23159.695 | 209 | A vs C | 8.444 | <0.01 |

| | | | | | | | |
|---|---|---|---|---|---|---|---|
| *ACPT* has $\omega_2$, others have $\omega_1$ | | | | | | | |
| D. The terminal branch of *Eubalaena glacialis* with pseudogenized *ACPT* has $\omega_2 = 1$, others have $\omega_1$ | $\omega_1$=0.116 | $\omega_2$=1 | 23160.587 | 208 | D vs C | 1.784 | 0.182 |
| **The terminal branch of *Balaenoptera bonaerensis*** | | | | | | | |
| A. All branches have one ω | 0.117 | | 23200.981 | 208 | | | |
| B. All branches have one ω = 1 | 1 | | 25437.712 | 207 | A vs B | 4473.462 | 0 |
| C. The terminal branch of *Balaenoptera bonaerensis* with pseudogenized *ACPT* has $\omega_2$, others have $\omega_1$ | $\omega_1$=0.116 | $\omega_2$=1.045 | 23190.135 | 209 | A vs C | 21.692 | <0.01 |
| D. The terminal branch of *Balaenoptera bonaerensis* with pseudogenized *ACPT* has $\omega_2 = 1$, others have $\omega_1$ | $\omega_1$=0.116 | $\omega_2$=1 | 23190.138 | 208 | D vs C | 0.006 | 0.938 |
| **The terminal branch of *Balaenoptera acutorostrata*** | | | | | | | |
| A. All branches have one ω | 0.117 | | 23178.238 | 208 | | | |
| B. All branches have one ω = 1 | 1 | | 25420.510 | 207 | A vs B | 4484.544 | 0 |
| C. The terminal branch of *Balaenoptera acutorostrata* with pseudogenized *ACPT* has $\omega_2$, others have $\omega_1$ | $\omega_1$=0.116 | $\omega_2$=0.613 | 23172.147 | 209 | A vs C | 12.182 | <0.01 |
| D. The terminal branch of *Balaenoptera acutorostrata* with pseudogenized *ACPT* has $\omega_2 = 1$, others have $\omega_1$ | $\omega_1$=0.116 | $\omega_2$=1 | 23172.621 | 208 | D vs C | 0.948 | 0.330 |
| **The terminal branch of *Balaenoptera borealis*** | | | | | | | |
| A. All branches have one ω | 0.118 | | 23231.146 | 208 | | | |
| B. All branches have one ω = 1 | 1 | | 25466.547 | 207 | A vs B | 4470.802 | 0 |
| C. The terminal branch of *Balaenoptera borealis* with | $\omega_1$=0.116 | $\omega_2$=0.902 | 23219.198 | 209 | A vs C | 23.896 | <0.01 |

| | | | | | | | |
|---|---|---|---|---|---|---|---|
| pseudogenized *ACPT* has $\omega_2$, others have $\omega_1$ | | | | | | | |
| D. The terminal branch of *Balaenoptera borealis* with pseudogenized *ACPT* has $\omega_2$ = 1, others have $\omega_1$ | $\omega_1$=0.116 | $\omega_2$=1 | 23219.224 | 208 | D vs C | 0.052 | 0.820 |
| **The stem group of Mysticeti** | | | | | | | |
| A. All branches have one $\omega$ | 0.121 | | 23747.298 | 224 | | | |
| B. All branches have one $\omega$ = 1 | 1 | | 25976.514 | 223 | A vs B | 4458.432 | 0 |
| C. The ancestral branch of stem mysticeti with *ACPT* has $\omega_2$, others have $\omega_1$ | $\omega_1$=0.120 | $\omega_2$=1.436 | 23741.059 | 225 | A vs C | 12.478 | <0.01 |
| D. The ancestral branch of stem mysticeti with *ACPT* has $\omega_2$ = 1, others have $\omega_1$ | $\omega_1$=0.116 | $\omega_2$=1 | 23741.164 | 224 | D vs C | 0.210 | 0.647 |
| **The crown group of Mysticeti** | | | | | | | |
| A. All branches have one $\omega$ | 0.121 | | 23747.298 | 224 | | | |
| B. All branches have one $\omega$ = 1 | 1 | | 25976.514 | 223 | A vs B | 4458.432 | 0 |
| C. The clade of crown mysticeti with *ACPT* has $\omega_2$, others have $\omega_1$ | $\omega_1$=0.116 | $\omega_2$=0.522 | 23720.907 | 225 | A vs C | 52.782 | <0.01 |
| D. The clade of crown mysticeti with *ACPT* has $\omega_2$ = 1, others have $\omega_1$ | $\omega_1$= | $\omega_2$=1 | 23725.565 | 224 | D vs C | 9.316 | <0.01 |
| **The terminal branch of *Kogia breviceps*** | | | | | | | |
| A. All branches have one $\omega$ | 0.117 | | 23212.116 | 208 | | | |
| B. All branches have one $\omega$ = 1 | 1 | | 25455.901 | 207 | A vs B | 4487.570 | 0 |
| C. The clade of *Kogia breviceps* with *ACPT* has $\omega_2$, others have $\omega_1$ | $\omega_1$=0.116 | $\omega_2$=0.581 | 23203.192 | 209 | A vs C | 17.848 | <0.01 |

| | | | | | | | |
|---|---|---|---|---|---|---|---|
| D. The clade of *Kogia breviceps* with *ACPT* has $\omega_2 = 1$, others have $\omega_1$ | $\omega_1$=0.116 | $\omega_2$=1 | 23204.052 | 208 | D vs C | 1.720 | 0.190 |
| **The terminal branch of *Dasypus novemcinctus*** | | | | | | | |
| A. All branches have one ω | 0.117 | | 23387.077 | 208 | | | |
| B. All branches have one ω = 1 | 1 | | 25653.621 | 207 | A vs B | 4533.088 | 0 |
| C. The terminal branch of *Dasypus novemcinctus* with pseudogenized *ACPT* has $\omega_2$, others have $\omega_1$ | $\omega_1$=0.116 | $\omega_2$=0.206 | 23384.542 | 209 | A vs C | 5.070 | 0.024 |
| D. The terminal branch of *Dasypus novemcinctus* with pseudogenized *ACPT* has $\omega_2 = 1$, others have $\omega_1$ | $\omega_1$=0.116 | $\omega_2$=1 | 23403.931 | 208 | D vs C | 38.778 | <0.01 |
| **The terminal branch of *Orycteropus afer*** | | | | | | | |
| A. All branches have one ω | 0.119 | | 23456.631 | 208 | | | |
| B. All branches have one ω = 1 | 1 | | 25695.648 | 207 | A vs B | 4478.034 | 0 |
| C. The terminal branch of *Orycteropus afer* with pseudogenized *ACPT* has $\omega_2$, others have $\omega_1$ | $\omega_1$=0.116 | $\omega_2$=0.414 | 23441.822 | 209 | A vs C | 29.618 | <0.01 |
| D. The terminal branch of *Orycteropus afer* with pseudogenized *ACPT* has $\omega_2 = 1$, others have $\omega_1$ | $\omega_1$=0.116 | $\omega_2$=1 | 23448.875 | 208 | D vs C | 14.106 | <0.01 |

### References:

Celine S, Khalid B, Jimmy L, Rémy D, Frédéric DE, J P Douzery Vincent, Ranwez: OrthoMaM v10: Scaling-up orthologous coding sequence and exon alignments with more than one hundred mammalian genomes. Molecular Biology and Evolution 2019, 36(4):861-862.

Gatesy J, Geisler JH, Chang J, Buell C, Berta A, Meredith RW, Springer MS, Mcgowen MR: A phylogenetic blueprint for a modern whale. Molecular Phylogenetics and Evolution 2013, 66(2):479-506.

Kuntner M, May- Collado LJ, Agnarsson I: Phylogeny and conservation priorities of afrotherian mammals (Afrotheria, Mammalia). Zoologica Scripta 2011, 40(1):1-15.

OrthoMaM: http://orthomam2.mbb.univ-montp2.fr/OrthoMaM_v10b10/.

Sergey N, Montoya-Burgos JI, Margulies EH, Jacques R, Bruno N, Antonarakis SE: Early history of mammals is elucidated with the ENCODE multiple species sequencing data. PLoS Genetics 2007, 3(1):e2.

Seymen F, Kim YJ, Lee YJ, Kang J, Kim TH, Choi H, Koruyucu M, Kasimoglu Y, Tuna EB, Gencay K: Recessive mutations in ACPT, encoding testicular acid phosphatase, cause hypoplastic amelogenesis imperfecta. American Journal of Human Genetics 2016, 99(5):1199-1205.

Smith CE, Whitehouse LL, Poulter JA, Brookes SJ, Day PF, Soldani F, Kirkham J, Inglehearn CF, Mighell AJ: Defects in the acid phosphatase ACPT cause recessive hypoplastic amelogenesis imperfecta. European Journal of Human Genetics 2017, 25(8):1015-1019.

Waddell PJ, Kishino H, Ota R: A Phylogenetic Foundation for Comparative Mammalian Genomics. Genome Informatics 2001, 12:141-154.

Xuming Z, Shixia X, Junxiao X, Bingyao C, Kaiya Z, Guang Y: Phylogenomic analysis resolves the interordinal relationships and rapid diversification of the laurasiatherian mammals. 2012, 61(1):150-164.
